# Neural mechanisms for the localization of externally generated tactile motion

**DOI:** 10.1101/2022.05.30.494073

**Authors:** Suma Chinta, Scott R. Pluta

## Abstract

During tactile localization, animals must differentiate stimuli caused by their own volitional movement from externally generated object motion. To determine a neural basis for this ability, we examined the mouse superior colliculus (SC), which contains multiple egocentric maps of sensorimotor space. By placing mice in a whisker-guided virtual reality, we discovered a rapidly adapting neural response that strongly preferred external over self-generated changes in tactile space. This transient response only emerged when external motion gained contact with a whisker, arguing that stimulus adaptation was whisker-specific. The accumulation of sensory evidence through active sensing and repetitions in external motion controlled the size of the transient response. Population-level firing rates among transiently responsive neurons accurately encoded the direction of external motion. These data reveal that stimulus-specific adaptation together with accumulating sensorimotor predictions in SC neurons enhance the localization of unexpected motion in the environment.

## Introduction

Much of our interaction with the world occurs through the volitional movement of our sensory organs. For example, primates move their eyes to visualize a scene, and rodents move their whiskers to explore nearby objects. To accurately localize objects, animals must differentiate the expected consequences of their actions, induced by active sensing, from real-world object motion. What neural mechanisms underlie this ability?

To address this question, we focused on the midbrain superior colliculus (SC), which contains multiple egocentric maps of sensorimotor space ^1–4^. Many SC mediated behaviors involve spatial processing, such as pursuing prey, escaping predators, or simply orienting towards the appropriate object ^5–11^. While visually tracking objects, the SC helps stabilize the visual field and adjust for discrepancies between the predicted and actual visual outcome of eye movements ^12,13^. Given this insight, we hypothesized that SC neurons are specialized for encoding environmental motion during active sensing. Such computations are likely critical for localizing moving objects such as prey ^14–17^.

While decades of research have outlined SC function during visuospatial processing, remarkably less is known for somatosensation, particularly during active touch ^18–21^. The mouse SC is known to respond to whisker stimuli, yet nearly all published work in this area has been performed under anesthesia ^4,22–25^. These experiments reveal that the SC contains a somatotopic map of whisker space, whereby individual neurons possess large (multi-whisker) receptive fields that are also sensitive to artificial whisking. Therefore, active whisking in the mouse SC provides a tractable and underutilized model for revealing the cellular and circuit mechanisms that support innate tactile-guided behaviors.

Sensory responses in the SC have primarily been characterized during sensor fixation, with evidence showing that differences in egocentric heading (eye position) modulate response magnitude ^18,19,21,26–28^. This approach reveals that sensory responses in the SC are modulated by extrasensory inputs. Additional work has shown that eye movements in darkness suppress SC spiking, arguing for the presence of a corollary discharge to suppress self-induced visual stimulation ^29,30^. The somatosensory whisker system offers a powerful model for investigating the integration of external and self-generated stimuli, since mice instinctively move their whiskers to palpate objects, and the precise dynamics of touch can be accurately measured using high-speed imaging and markerless tracking ^31,32^.

To reveal the neural mechanisms for the active localization of external motion, we designed a tactile virtual reality that simulates whisker-guided navigation. While running on a treadmill, mice rhythmically touched a cylindrical surface that rotated at the same velocity as their locomotion. This created a tactile flow-field that simulates running along a wall, in a manner similar to visual flow ^33–35^. Periodically, the center axis of the flow-field translated horizontally along the mouse’s whisking field. A transient neural response only emerged when surface movements entered a location that gained or lost contact with a whisker. Self-generated gains in whisker contact that occurred independent of surface movement generated relatively weak tactile responses. The size of the tactile response was controlled by stimulus expectations, and the direction of surface movement was accurately encoded in neuronal firing rates.

## Results

### Simulating whisker-guided virtual reality

We designed a closed-loop tactile system that simulates whisker-guided exploration in mice. During the experiment, a head-fixed running mouse voluntarily whisked against a nearby cylindrical surface that rotated at the same velocity as its locomotion (**Fig. 1A, left**). A high-speed (500fps) infrared camera and digital encoders recorded whisker kinematics, surface movements, and running speed while an opaque object and white noise obscured visual and auditory cues. After the mouse initiated a trial by running a certain distance, the surface would either translate rostrally, caudally or remain at the starting location with equal probability. After staying at the rostral/caudal location for an equivalent locomotor distance (if applicable), the surface would then return to the center location. While running and touching the surface, mice innately adapted the position of their whiskers to track changes in surface location (**Fig. 1A, right**). Since translation timing was determined by locomotor distance, variation in locomotion speed made translation onset unpredictable (**Fig. 1B**). As the surface moved into a new location, the firing rate of most neurons rapidly increased while others decreased, as shown in two example units (**Fig. 1C**). A 3-shank, 128-channel silicon probe recorded neural activity simultaneously across the intermediate and deep layers of the lateral SC, approximately 300 – 1000 microns below its surface (**Fig. 1D**, 12 mice, 873 neurons). About two-thirds of all recorded neurons displayed a significant response to surface movement (67 ± 6%, 12 mice, 578 neurons, α < 0.05, 1-way ANOVA).

**Figure 1.**
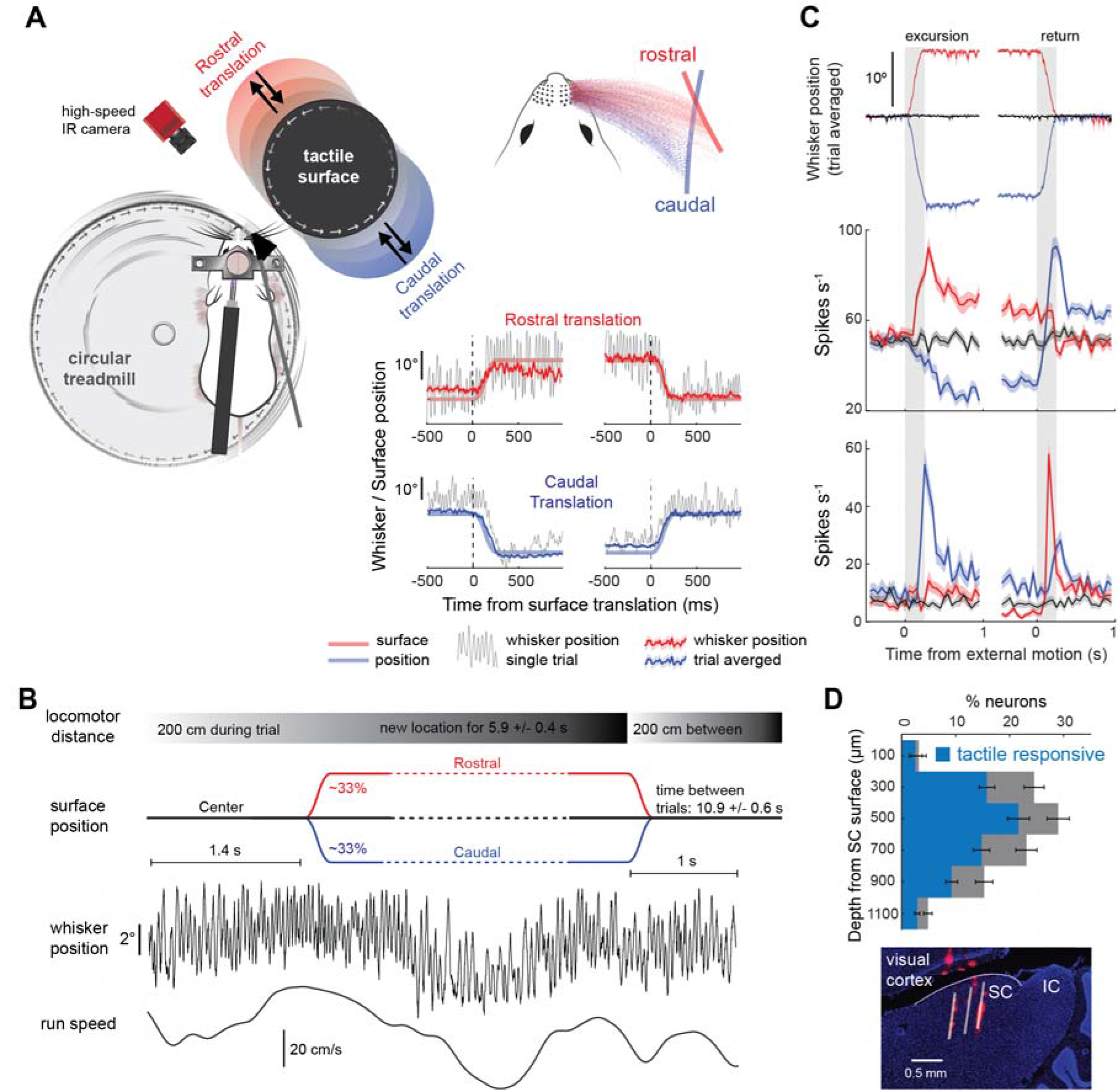
Tactile and neural dynamics during whisker guided virtual reality. **A**) Schematic of whisker guided virtual navigation, illustrating a head-fixed mouse locomoting on a treadmill and whisking against a surface. After trial initiation, there was an equal probability for the tactile surface to translate into either rostral (in red) or caudal (in blue) whisker space or stay at the center position (in black). Upper right: single-whisker location density while the surface was at the rostral or caudal location. Bottom right: whisker position (mean ± s.e.m) and surface location (transparent line) during rostral (top in red) and caudal (bottom in blue) translations. The first half is surface excursion while second half is the return to the center position. **B**) Trial structure aligning surface position, whisker position, and locomotion speed. **C)** Trial-averaged whisker position and firing rate of two example SC neurons during each type of surface translation (50ms bins). **D)** Top: Percentage of tactile responsive SC neurons (blue) as a function of depth from the SC surface (12 mice, 873 neurons). Bottom: dye labeling of the electrode shanks in the intermediate and deep layers of the lateral SC.

### Surface movements that gain or lose whisker contact elicit transient and sustained changes in firing rate

To reveal the stimulus variables driving SC responses, we examined the dynamics of whisker touch during surface movement. We performed a combination of computerized whisker kinematic analysis and manual inspection of high-speed video. We discovered that surface movements elicited large changes in firing rate only when they entered a location that gained (GoW) or lost (LoW) contact with a whisker (**Fig. 2A, B, Suppl. Fig. 1A**). The addition or subtraction of a single whisker from the surface was sufficient for this effect. Most neurons preferentially responded to whisker addition (GoW, 55 ± 4%, 8 mice, 385 neurons), while only a small percentage of neurons preferred surface movements that engaged the same whiskers at both locations (15 ± 3%, 8 mice, 385 neurons, **Supp. Fig. 1A**). Gains in whisker contact with the surface elicited transient and/or sustained changes in neuronal firing rate (**Fig. 2A, C top**). Transient responses typically returned to baseline within 300 ms following their onset, while sustained responses exhibited firing rates different than baseline for the entire duration that the surfaced contacted the new whisker(s) (see methods). In most cases, the transient response was larger than the sustained response, yet there was almost continuous variation in their relative magnitudes across the population (**Fig. 2C, bottom**). To confirm that these responses dynamics were whisker-mediated, we repeated the experiment after trimming off the whiskers and found that nearly all neurons became unresponsive to surface movement (59 ± 4% pre-trimming vs. 6 ± 1% post-trimming, 156 neurons, 2 mice, 1-way ANOVA, α < 0.05, **Supp. Fig. 1B**). To test if adaptation of the transient response was whisker specific, we picked trials where two consecutive surface movements gained contact with different whiskers. With each consecutive gain in whisker contact, a transient response emerged from the adapted state (**Supp. Fig 1C**).

**Figure 2.**
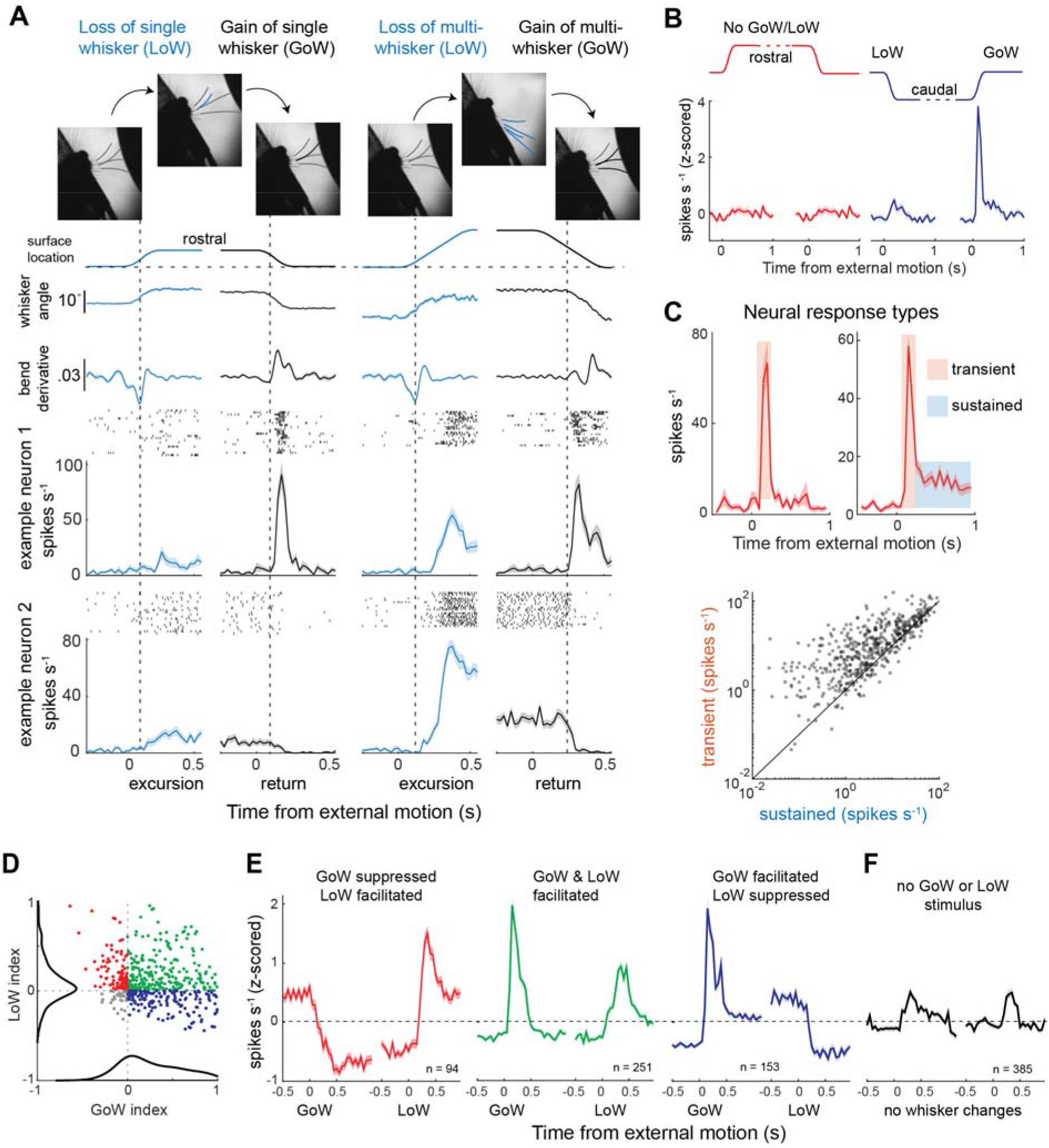
Surface movements that gained or lost contact with whiskers elicited transient and sustained changes in spike rate. **A**) Surface location, whisker angle, whisker curvature derivative, and the firing rates of two example neurons during externally generated gains (black) and losses (blue) in whisker touch. The vertical dashed line marks the estimated moment of gain/loss in contact. **B**) Population firing rates from an example mouse during surface movements. A gain/loss in whisker contact only occurred during movement through caudal space (46 neurons). **C**) Top, two example neurons illustrating the distinct transient and sustained response components evoked during surface translation. Bottom, scatter plot comparing the transient and sustained firing rates of SC neurons (10 mice, 529 neurons). **D**) Scatter plot comparing the GoW and LoW modulation of each neuron (10 mice, 529 neurons). **E**) Population averaged firing rates of neurons grouped by the sign (+/-) of their responses to GoW and LoW (8 mice, 385 neurons). **F)** Population average firing rate of neurons response to no GoW or LoW external surface movement. All population average firing rates are binned at 50ms.

To understand the receptive fields of SC neurons in the framework of whisker space, we quantified each neuron’s response to GoW and LoW stimulation (10 mice, 529 neurons, **Fig. 2D**). Almost half of all neurons increased their firing rate for both GoW and LoW (251/529 neurons) stimuli, while the remaining neurons increased their firing rate for one stimulus type yet decreased it for the other. To illustrate these effects, we plotted the population averaged firing rates of each response type relative to the onset of surface movement (**Fig. 2E**). Overall, these data demonstrate that SC neurons were highly sensitive to gains/losses in whisker contact, while showing relatively weak changes in firing rate during surface movements that sustained contact with the same whiskers (**Fig. 2F**).

### SC neurons strongly prefer externally generated gains in whisker contact

We hypothesized that the GoW response was more sensitive to external rather than self-generated gains in whisker contact. To test this hypothesis, we modified our experiment so that the surface periodically moved between rostral whisking space and a more distant location entirely outside the whisking field. With the surface in rostral space, mice voluntarily lost and gained contact (self-GoW) with the surface as they periodically (6.8 ± 0.6 second interval) transitioned between resting (no running or touch) and locomotive touch (**Fig. 3A**). Mice had 1 to 5 intact whiskers contacting the surface in these experiments. Using this approach, we were able to calculate neuronal firing rates relative to the onset of self-GoW as well as externally generated gains in whisker contact (external-GoW).

**Figure 3.**
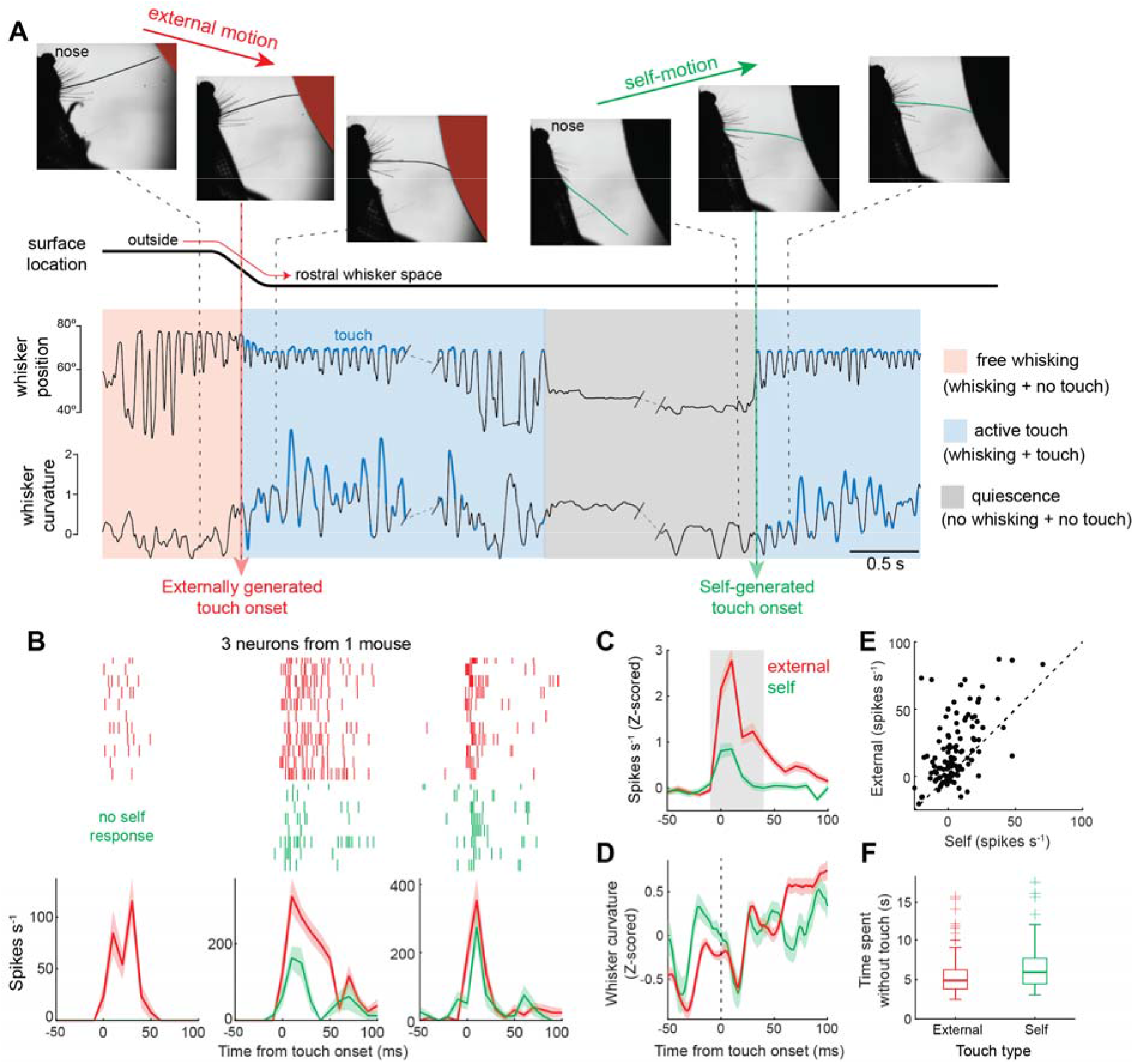
Externally generated gains in contact preferentially drive the transient response. **A**) Example trial illustrating the designation of external- and self-generated touch. Initially, the mouse freely whisked in air. After the mouse ran a set distance, the surface moved into its whisking field and contacted the whisker to create externally generated touch. After a period of whisking against the surface, the mouse voluntarily stopped before eventually resuming to create self-generated touch. **B**) Three neurons from one mouse comparing responses to external and self-generated touch. Rasters and histograms of spike timing aligned to touch onset. Only the first external and self-generated touch from each trial were used. **C**) Population-averaged, z-scored firing rates aligned to the onset of external and self-GoW (4 mice, 139 neurons). Firing rates for external and self-GoW plotted in E are extracted from the gray window. **D**) Population averaged, z-scored whisker curvature aligned to the onset of external and self-GoW. **E**) Scatter plot comparing change in neuronal firing rates for external and self-generated touch (4 mice, 139 neurons, p < 0.001, Wilcoxon signed rank test). Firing rates are subtracted with their corresponding baselines. **F**) Boxplots comparing the time mice spent without any touch before an external- or self-GoW stimulus occurred (p = 0.04, Mann Whitney U-test). All values are mean ± s.e.m.

In individual mice, neurons displayed a range of stimulus sensitivities (**Fig. 3B**). Some neurons only responded to external-GoW stimuli, while most neurons responded to both stimulus types but with a clear preference for external-GoW (**Fig. 3B, C**). A minority of neurons responded equally to external- and self-GoW stimuli (**Fig. 3E**). Therefore, population level firing rates were significantly greater during external-GoW stimulation (p < 0.001, Wilcoxon signed rank test, 4 mice, 139 neurons, **Fig. 3C, E**). The curvature of the whisker during external and self-GoW was nearly identical, indicating that differences in stimulus strength are unlikely the cause of this effect (**Fig. 3D, Suppl. Fig. 2A**). To rule out adaptation as the mechanism for the larger external-GoW response, we compared the time mice spent without touch preceding each GoW event. Our data indicate that adaptation cannot explain GoW response differences, as the time spent without touch before GoW stimulation was similar in both conditions (Mann-Whitney, p = 0.04, **Fig. 3F, Suppl. Fig. 2B**). Therefore, the larger external-GoW response cannot be explained by a slower stimulus repetition rate.

### Self-generated sensory experience controls transient response magnitude

Although stimulus strength and repetition rate for external- and self-GoW stimuli were the same, the locomotive states were not. In the external-GoW condition, the animal was already running, yet in the self-GoW condition, the animal was transitioning from quiescence to locomotion. Using a novel approach where both touch events started from quiescence, we controlled for differences in locomotion speed and any potential differences in touch quality (**Fig. 4**).

**Figure 4.**
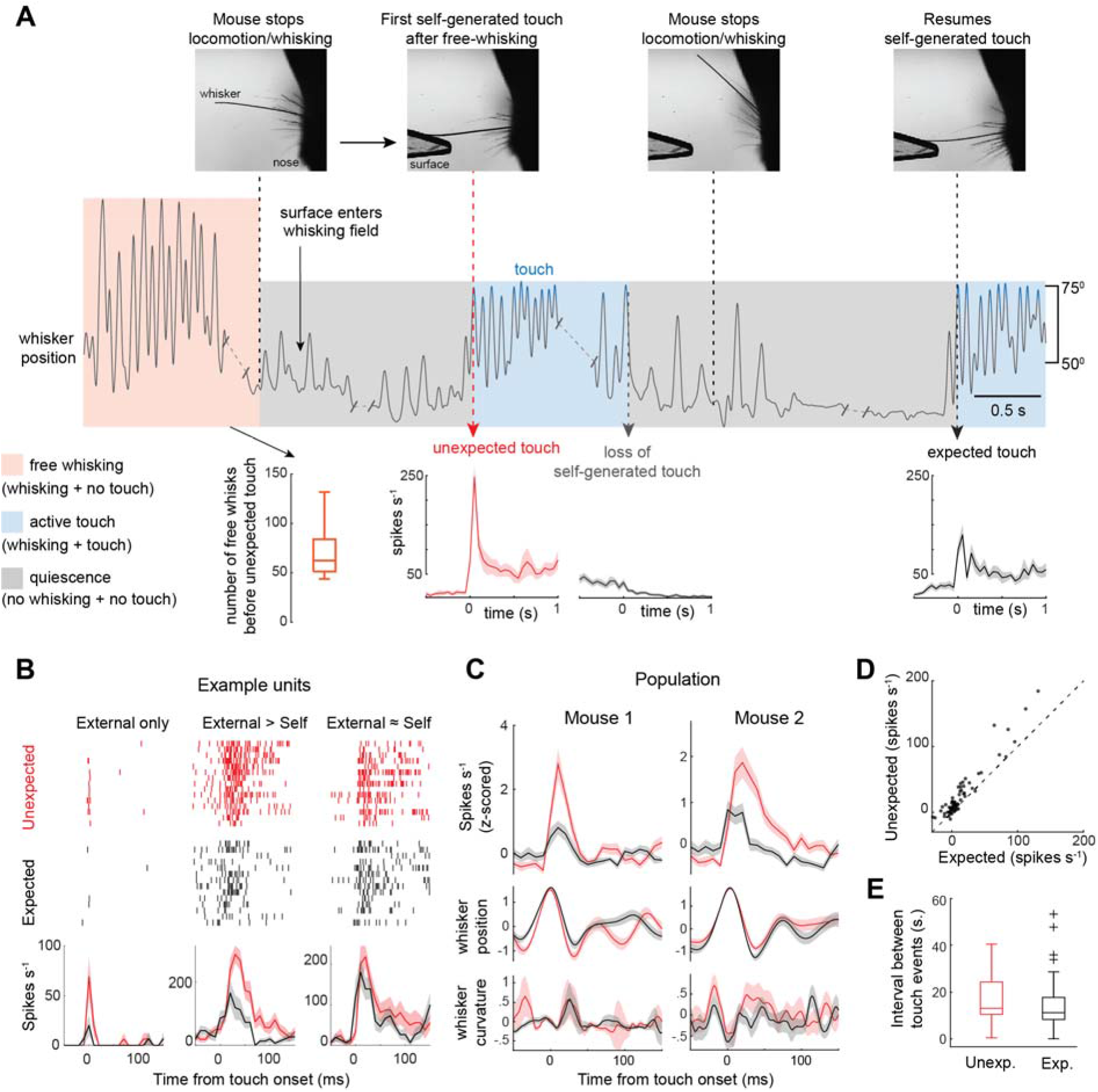
Self-generated stimulus expectations control the size of the transient response. **A)** Schematic illustrating the experimental design. At the start of the trial, the mouse is running and whisking freely in air (see boxplot for quantification). After the mouse crossed a threshold locomotion distance and then stopped running, a touch surface entered its whisking field. The mouse voluntarily resumed whisking and generated an unexpected touch event. The mouse then voluntarily stopped running and whisking before it resumed active touch. After running and touching the surface for a set distance, the surface retracted, and the trial restarted. *Bottom*, histogram of firing rates for one neuron during expected and unexpected touch. **B)** Rasters and histograms of spike timing for three neurons from one mouse during expected and unexpected touch. **C)** Population averaged neuronal firing rates and whisking variables aligned to the onset of touch (2 mice, 74 neurons). **D)** Scatter plot comparing change in neuronal firing rates during expected and unexpected touch (2 mice, 74 neurons, p = 0.004, Wilcoxon signed rank test). **E)** Box plot comparing the time mice spent without any touch before an unexpected and expected touch event occurred (Mann Whitney U-test, p = 0.17).

Mice had one whisker contacting the surface in this experiment. A trial was initiated by the animal running and freely whisking in air. After the mouse reached a threshold distance on the treadmill, and then decided to stop running and whisking, an object was quickly (150 ms) extended into its whisking field (**Fig. 4A**, see boxplot). Moments later, the animal voluntarily resumed locomotive whisking and contacted the surface, causing an unexpected touch event. The object stayed in the whisking field for an extended period, allowing the mouse to periodically accumulate evidence about its presence by pausing and resuming active touch. After each pause in locomotion and whisking, the first touch in the next locomotive whisking bout generated an expected touch event. After the mouse ran and touched the surface for a set treadmill distance, the surface retracted, and the trial restarted with the free whisking and locomotion condition. On average, the first touch response that followed free whisking (unexpected) was significantly greater than the first touch response that followed a bout of active touch (expected, **Fig. 4B, C,** p = 0.004, n = 74, signed rank test). Importantly, touch kinematics (whisker position and whisker curvature) were identical between both touch conditions (**Fig. 4C**). Neurons with larger tactile responses were more strongly modulated by stimulus expectation (**Fig. 4D**). Stimulus repetition rate was the same between the touch conditions (**Fig. 4E,** p = 0.17, t-test). Therefore, stimulus expectations (whisking in air vs. repeated surface contact) generated by active sensing controlled the magnitude of the transient response, in agreement with our data in Figure 3.

### Repetitions in external motion weaken the transient response

Next, we sought to determine if responses to external-GoW were static or habituated over time. To do so, we analyzed the effect of stimulus repetition on response magnitude. For each experiment, we selected a surface movement with GoW stimulation (**Fig. 5A**). To ensure that any effects we observed were not caused by changes in tactile behavior, we analyzed the position of the whisker as a function of stimulus repetition. Across the population (8 mice), there was no notable change during the transient response period (**Fig. 5B**). However, we discovered that the magnitude of the transient response linearly decayed with increasing GoW repetitions, while the sustained response was stable, as shown in two example neurons (**Fig. 5C, D**). This transient-specific habituation was consistent across the population (**Fig. 5E**, 223 transient neurons, 8 mice, p = 2e^-12^; 98 sustained neurons, p = 0.64, t-test). The stability of spike waveform amplitude in our recordings indicates that a progressive decrease in spike detection during sorting cannot explain these effects (**Fig. 5F, 5C insets**). Habituation of the transient response was unlikely caused by low-level sensory adaptation, since the average interval between each GoW stimulus was around one minute (67 ± 5 sec, 8 mice), and GoW stimuli were randomly interleaved with other non-GoW surface movements. Therefore, SC neurons appear to steadily habituate to repeated tactile stimulation occurring over the course of many minutes, providing further evidence for the role of sensory experience in regulating the transient response. The same transient-specific habituation was observed in our other approach detailed in Figure 4 (**Suppl. Fig. 3**).

**Figure 5.**
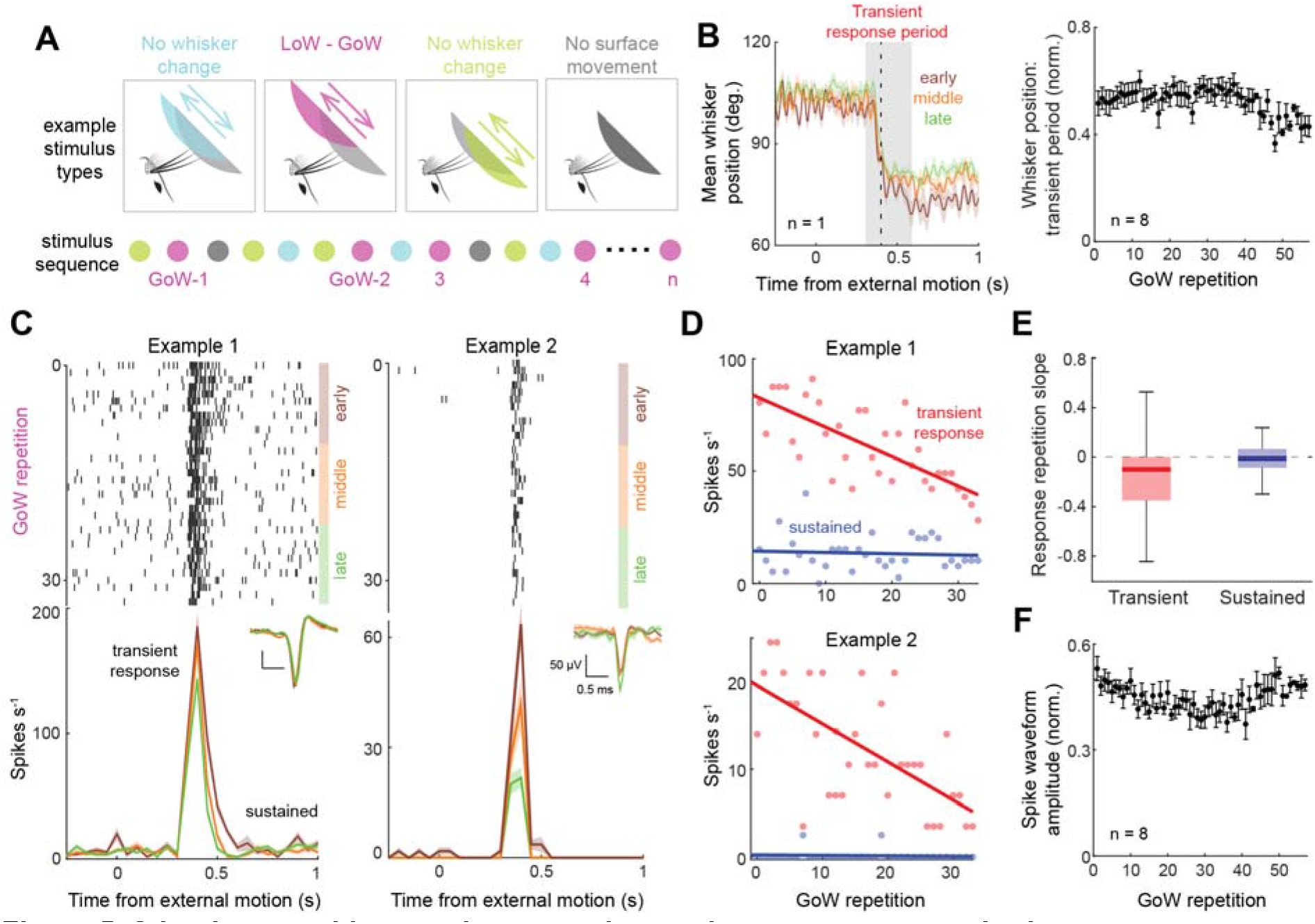
Stimulus repetition number controls transient response magnitude. **A**) Schematic illustrating the different stimulus types and how they are randomly repeated over the course of the experiment. The return motion of the 2^nd^ stimulus which is a GoW is selected for analysis in C. **B**) Left, trial-averaged whisker position relative to surface movement for early (first 1/3^rd^), middle, and late (last 1/3^rd^) trials of GoW stimulation (one mouse). Right, trial-averaged whisker position calculated during the transient response period and plotted as a function of stimulus repetition (8 mice). **C)** Rasters and histograms of spiking in two example neurons divided by early, middle, and late periods. Inset, spike waveforms during the same periods (color coded). **D)** Magnitude of the transient and sustained responses as a function of stimulus repetition in same two example neurons in **C**. Transient and sustained firing rates were obtained from transient and sustained windows described in methods. **E)** Population slopes of the linear regression of transient (8 mice, 223 neurons, p < 0.001, t-test) and sustained (8 mice, 98 neurons, p = 0.64, t-test) responses as a function of stimulus repetition. **F)** Normalized spike waveform amplitude as a function of stimulus repetition (8 mice).

### Neurons in the superior colliculus encode surface location

To assess the function of external and self-generated tactile responses in the SC, we tested the surface localization accuracy of population-level SC activity over time. Using population decoding (see methods), we discovered that SC activity accurately classified surface location (center, rostral, or caudal), with the highest accuracy during the movement period, as shown in one example mouse (**Fig. 6A**, 99% accuracy, 76 neurons). Before the start of the excursion, classification accuracy was at chance, indicating that neural activity could not predict the upcoming surface movement. After excursion into rostral or caudal space, classification accuracy for sustained neurons was high for as long as the surface remained at that location, revealing a stable self-generated representation of location in this subset of neurons (**Fig. 6B**, 30 neurons, 1 mouse). After the surface returned to the center position, stimulus classification remained above chance, revealing persistent information about the previous location of the object. In contrast, classifier accuracy for transient neurons was high only during surface movement, due to their rapid adaptation rates (46 neurons, 1 mouse). By segregating experiments by the number of surface movements that gained/lost contact with a whisker, we found that classifier accuracy was highest during GoW/LoW translations (**Fig. 6C, D**, GoW/LoW in all movements: 94 ± 5% accuracy, 2 mice; GoW/LoW in half: 71 ± 7%, 6 mice; GoW/LoW in no movements: 56 ± 3%, 4 mice). This trend was generally consistent for both sustained and transient neurons (**Fig. 6E**), but transient neurons were least informative about surface movements that did not gain or lose whisker contact. Since classifier accuracy was best at distinguishing locations that interacted with different whiskers, a somatotopic map of whisker space in the SC likely facilitates object localization. We found evidence for such map in one of our recordings where the rostral and caudal stimuli gained contact with different whiskers (**Suppl. Fig. 4**).

**Figure 6.**
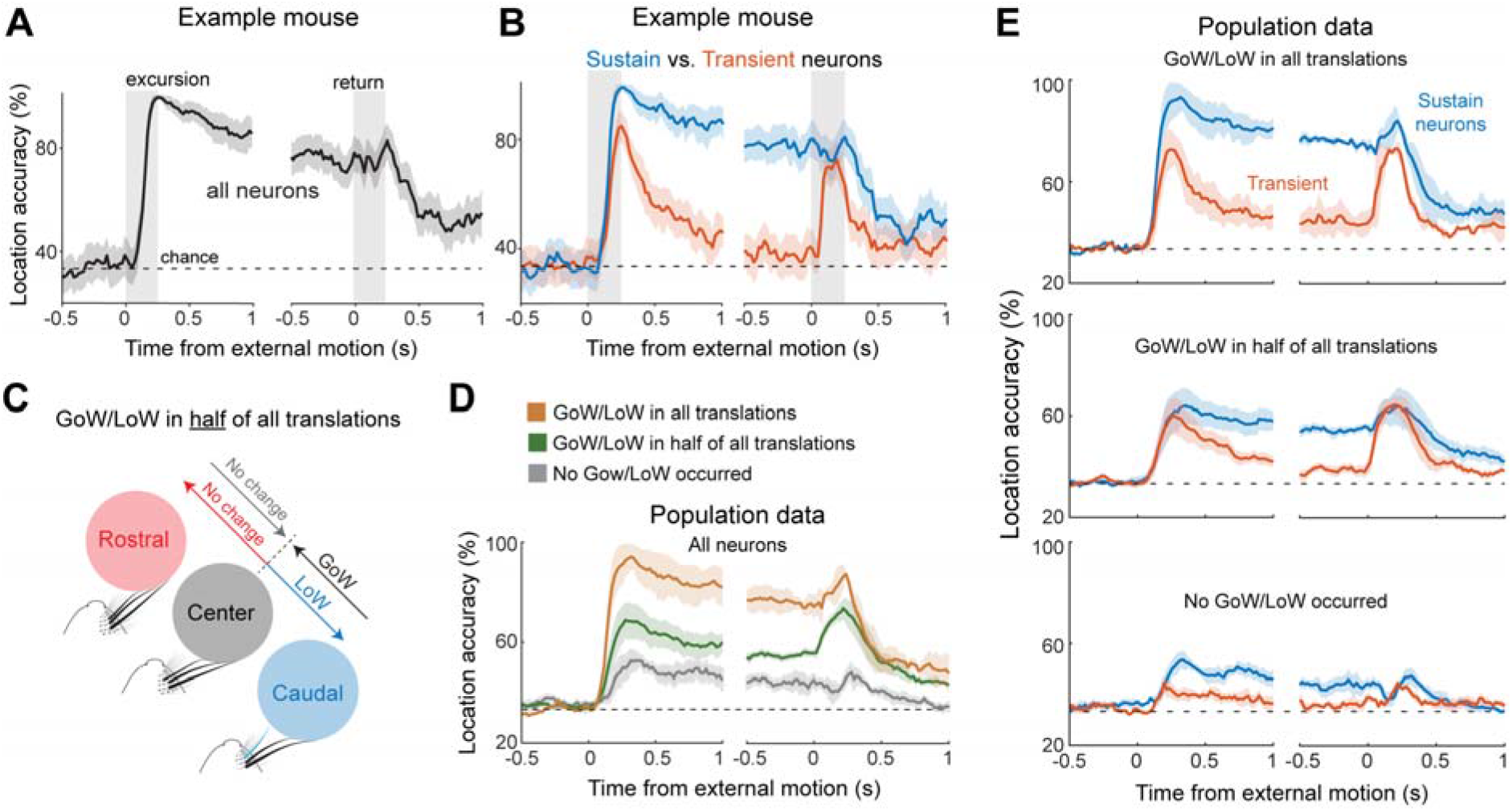
Neural decoding of surface location. **A)** Location decoding accuracy using all the tactile responsive neurons recorded in a single mouse (75 neurons). **B)** Decoding accuracy in the same mouse with sustained (n = 29) and transient (n =34) responsive neurons separated. **C)** Diagram illustrating an experiment where half of all the surface translations caused a gain/loss in whisker contact. **D)** Decoding accuracy averaged across animals with the same number of gains/losses in whisker contact (GoW/LoW in all movements: 2 mice, 81 neurons; GoW/LoW in half of all movements: 6 mice, 262 neurons; Zero GoW/LoW movements: 4 mice, 149 neurons). **E)** Decoding accuracy averaged across animals according to neuron type (sustain/transient) and changes in whisker contact (12 mice, 492 neurons).

## Discussion

Active sensing requires the brain to contextualize incoming sensory stimuli with outgoing motor commands. To localize objects, animals must recognize if changes in egocentric space originate from externally or internally generated movement ^36^. By simulating whisker-guided exploration, we discovered neurons in the mouse SC that are specialized for locating unexpected, externally generated changes in tactile space. We identified a rapidly adapting transient response, which strongly preferred externally initiated touch and that habituated with stimulus repetition. Transient responses have been observed in the SC during visual stimulation ^37–39^ and passive, rhythmic deflection of the whiskers ^25^. We observed a diversity of stimulus selectivity across the population of recorded neurons, with some neurons only briefly responding to externally generated touch and other neurons sustaining an equally strong response to both external and self-generated touch. Interestingly, self-generated sensory experience appeared to be a primary factor controlling transient response magnitude.

Both the barrel cortex and brainstem provide monosynaptic sensory drive to SC neurons ^22–24,40,41^. While the circuit mechanism for rapid sensory adaption is unknown^42^, some layer 5 neurons in the barrel cortex also show rapid sensory adaption during surface whisking^33^. Brainstem derived synaptic potentials in SC neurons also rapidly adapt during 20 Hz stimulation, which is similar to the natural whisk frequency of mice^22^. Therefore, sensory adaptation in the SC could stem from local and/or long-range circuits. Relating the adaptation rate of SC neurons to their downstream targets could provide valuable insight into the mechanisms supporting SC-mediated movements and attention^10,43^.

How motor and associative circuits in the SC contextualize somatosensory input remains unknown. An efferent copy of whisker protraction, possibly involving the cerebellum, motor cortex and brainstem, could be an important mechanism for regulating the transient response ^13,44–48^. In our study, stimulus expectations built by active whisking shape transient response magnitude. Therefore, the magnitude of the transient response reflects the degree of mismatch between the predicted and actual outcome of sensor movement, thereby providing a neural mechanism to identify and correct for errors associated with sensory-guided movement^13^.

Value-coded input to the SC could shape the transient response to generate a neural mechanism for favoring novel or rewarding environmental cues ^49–55^. Importantly, mice in our study behaved voluntarily and were not seeking reward. In this free-form context, transient response habituation may reflect a central mechanism for ignoring stimuli of diminishing novelty ^38,56^. Whether this mechanism relies on changes in inhibitory signaling from the substantia nigra, the activity of local SC interneurons, or other upstream circuits remains unknown ^6,52,57^ Habituation to repeated visual stimulation has been observed previously in primates and rodents ^38,58–60^, yet our study is the first to demonstrate slower, accumulative effects of stimulus repetition in the SC during active sensing. Since our stimuli repeated at long, one-minute intervals over the course of an hour, SC neurons in our experiment were storing and updating stimulus expectations for an extended period.

Neuronal selectivity for externally generated stimulation has been observed in other sensory modalities and brain areas. Neurons in the monkey cerebellum selectively respond to external but not self-generated vestibular accelerations ^61^. Neurons in the monkey visual cortex display a unique window of suppression only when voluntary eye-movements create visual stimulation ^62^. Neurons in the superficial visual cortex of mice selectively respond to mismatches between visual flow and locomotion speed ^63,64^. Neurons in the auditory brainstem cancel self-generated sounds associated with licking ^45^, and neurons in the hindbrain of weakly electric fish cancel self-generated electric signals ^65^. We revealed a class of neurons in the superior colliculus that preferentially respond to externally generated tactile stimuli. Since contact with a new whisker was necessary for this transient response to emerge, the cancellation of self-generated touch is specific to individual whiskers.

Transient responses may facilitate object localization. We show that their population level firing rates briefly but accurately encode the location of external motion. The net effect of this coding brevity is an accurate representation motion direction, but not its lasting location. Decoding external motion was most accurate between movements that engaged different whiskers. Therefore, the somatotopic map of whisker space in the SC may enable a population level representation of object movement through different whiskers in the pad. The precise timing of spikes generated by these gains in whisker contact, as demonstrated in our study, could enable the rapid orienting movements associated with prey capture ^17,66^.

To localize objects during active touch, animals must differentiate self-from externally generated changes in tactile space. Such computations are pervasive across different sensory modalities and brain areas involved in sensorimotor processing. In the SC, response selectivity for unexpected, externally generated touch is controlled by rapid stimulus adaptation and the accumulation of external and self-generated stimulus expectations. These computations require single neurons to contextualize stimulus information over milliseconds and hours. It is perhaps surprising that single neurons in an ancient midbrain structure can multiplex this level of information. Given the diversity and complexity of circuits that impinge upon the mammalian SC, this may also be a reasonable expectation. How the SC interprets each of its diverse inputs to construct the egocentric maps that guide behavior remains an exciting and important topic.

## Methods

### Experimental Model and Subject Details

Mice of a CD-1 background of both sexes, between the ages of 9-15 weeks were used for all experiments. The Purdue Institutional Animal Care and Use Committee (IACUC, 1801001676A004) and the Laboratory Animal Program (LAP) approved all procedures. Mice were socially house with five or less per cage and maintained in reverse light-dark cycle (12:12 hr.). All experiments were conducted during the animal’s subjective night.

### In-vivo Electrophysiology

Prior to behavioral conditioning, a custom aluminum headplate was attached to each mouse to enable head fixation. Briefly, animals were anesthetized under 5% isoflurane and maintained at ~3% while monitoring body temperature and respiratory rate throughout the procedure. Artificial tears ointment was applied to keep the eyes hydrated. The skin and fur over the skull were disinfected with 70% ethanol followed by betadine and incised using sterilized surgical instruments. A tissue adhesive (Liquivet) and dental cement (Metabond) was applied over the skull and wound margins. The headplate was attached to the skull with dental cement. Lastly buprenorphine (mg/kg) was administered as a postoperative analgesic. Two days after headplate implantation, mice were placed on a circular treadmill for 1 hour per day for up to 10 days, or until they learned to run freely at a steady pace.

On the day of the electrophysiology recording, mice were briefly (15 – 20 minutes) anesthetized to perform a craniotomy over the SC. A 1mm diameter craniotomy was made using a biopsy punch (Robbins Instruments) and then covered with a silicone gel (Dowsil). Several hours later, mice were placed in the experimental rig and a three-shank custom probe (Neuronexus) of 128-channels was lowered into the brain using a micromanipulator (NewScale). After exiting the ventral surface of the cortex, as evident by a loss of spiking, the electrode was lowered at a rate of 75μm/min while constantly searching for activity driven by flashes of light. The onset of visual activity was used to mark the depth of the SC surface (~1000μm below cortical surface). The electrode was further lowered into the intermediate and deep layers, where manual whisker deflections caused strong neuronal responses. The receptive field of neurons were mapped by manually deflecting individual whiskers and locating the primary drivers of neuronal activity. Whiskers that did not elicit detectable activity were trimmed. Recordings were targeted towards the C-row and macrovibrissae. If the electrode penetration missed the target, it was removed and re-inserted based on the coordinates of somatotopic space. In most experiments, mice had 3 – 5 intact whiskers contacting the surface, which spanned one or two rows. In other cases, mice had one or two whiskers intact.

### Whisker guided virtual navigation

Head-fixed mice ran at their own volition (23.7 ± 1.6 cm/s, 12 mice) with concomitant rhythmic whisking (19 ± 1 Hz, 10 mice). All data presented was collected during locomotion and whisking, unless otherwise stated. The circular treadmill was attached to a digital encoder that controlled the rotation of a tactile surface in a closed-loop. A trial began after 200 cm of locomotion on the treadmill to ensure the mice were actively sensing the surface. Shortly (1.4 sec.) after whisker imaging began, the surface translated 1 cm linear distance into rostral or caudal space or remained at the center location. Each outcome was randomly chosen with equal (33.33%) probability. In six mice, an additional location was added that was entirely outside the whisker field, giving each surface movement a 25% probability. While at the rostral/caudal position, mice had to run an additional 200 cm before the surface would return to the center. An opaque flag over the eye and white background noise obscured visual and auditory cues, respectively.

In two mice, the tactile surface was replaced with a pneumatically controlled rectangular surface. Prior to the experiment, all whiskers were trimmed except the principal whisker (B1 or C1). A trial began after the mouse ran 200 cm on the treadmill while freely whisking in air with no surface in the whisking field. After the mouse crossed the distance threshold and stopped running, the touch surface was extended into the whisking field. The mouse voluntarily resumed whisking and generated an unexpected touch event. The mouse then voluntarily stopped whisking for a period before resuming to touch the same surface and generate an expected touch event. After running 600 cm with the surface present, the surface retracted, and the trial restarted.

### Whisker Tracking & kinematics

Whiskers were tracked at 500fps during the trial. A high-speed infrared camera (Photonfocus DR1) and a mirror angled at 45 degrees captured whisker motion under IR illumination. Videos were synchronized with neural data via external triggers generated by a National Instruments card and recorded on an Intan 512 controller. DeepLabCut was used to label the whisker(s) and track their movement ^31^. Four evenly spaced points on each whisker were labeled.

Approximately 150 frames were manually labeled from each experiment spanning all surface positions. The DLC neural network was trained for at least 200k iterations and the final labels were manually checked for accuracy. Whisker position/angle was calculated for each label on each whisker with reference to a user defined point on the face relative to the frame’s vertical axis. Whisker angle was bandpass filtered (1-30Hz, fdesign.bandpass order 4, MATLAB). Whisker bend/curvature was calculated from the three distal labels on each whisker using Menger curvature. The whisker curvature derivative was calculated as the local slope with cubic approximation in a moving window of 100ms.

### Spike Sorting

Spikes sorting was performed using the Kilosort2 ^67^ and manually curated using Phy2 gui (https://github.com/cortex-lab/phy). Spike clusters were considered single units based on their spike waveform, location on the electrode, and cross-correlogram of spike times. Single units were used for all analyses in the paper.

#### Statistical Analyses

##### Surface movement statistical classifier

To classify the location of the tactile surface from population-level SC activity, we applied the Neural Decoding Toolbox ^68^ using a support vector machine (SVM). The classifier predicted surface location (rostral, caudal, center) throughout the duration of the trial. Only neurons with a significant response to surface translation were used. Spike data was first Z-scored to prevent high firing rate neurons from having a disproportionate influence on classification. The spike rate of every neuron was calculated using 150ms bins and surface location was predicted every 20ms to plot classification accuracies over time. A 10-fold cross validation was performed along with 50 resample runs to calculate a robust estimate of classification accuracies. The classification accuracy was measured based on the zero-one loss function.

##### Modulation Indices

The gain and loss of whisker modulation indices (GoW/LoW indices) were calculated as the difference divided by the sum of spike rate averages that were calculated 500 ms before and after the start of surface movement.

##### Transient and sustained neurons classification

Surface movement was defined as a GoW/LoW stimulus when at least one whisker gained or lost contact. Since changes in whisker contact (GoW/LoW) occurred at slight time differences relative to the onset of surface movement across mice, we selected a custom transient window for each mouse based on the population average firing rates for every GoW/LoW translation. The peak firing rate of the transient response was identified by eye and the troughs around the peaks were manually marked as the start and the end of the transient window. We verified this window onset with analysis of whisker kinematics (whisker curvature and position). The firing rate of the sustained response was calculated in a 300 ms window that started one second after the transient window ended. We performed a one-way ANOVA (MATLAB anova1) on the firing rates calculated in the baseline (300 ms pre-movement), transient, and sustained windows for every neuron. A Tukey post hoc test (multcompare, MATLAB) was used to test for significant (α < 0.05) differences, correcting for multiple comparisons. Neurons which had significantly different baseline and sustained firing rates were classified as sustained neurons. Neurons with a transient window firing rate that was greater than both their baseline and sustain window firing rates were classified as transient neurons. Neurons which had a significant transient and/or sustained response were considered tactile responsive and used for analysis.

##### Self-GoW and external-GoW (Figure 3)

Self-GoW and external-GoW firing rates were calculated in a 50ms window following touch onset. Self-GoW was calculated as a difference between mean firing rates during self-GoW and self-motion. Self-motion occurred when the animal transitioned from resting to locomotion but free-whisking in air. The external-GoW response was calculated as the mean firing rate in the touch window minus the pre-touch (free-whisking) window.

##### Expected and Unexpected touch (Figure 4)

Expected and Unexpected touch firing rates were calculated by subtracting pre-touch firing rates from post touch in a 50 ms window.

##### Stimulus repetition

To obtain repetition slopes, we fit firing rates of TR neurons in the TR window and SR neurons in the SR window with a linear regression model using the MATLAB fitlm function. We only used neurons that did not drift. Neuronal drift was determined by testing if the baseline firing rate during the first 10 trials was different from the last 10 trials using a Wilcoxon signed rank test. Spike amplitudes were obtained from ‘amplitudes.npy’ output from Phy GUI. Normalized whisker position across repetitions were obtained as the mean of the normalized whisker position in the transient response window for each repetition. Spike waveform amplitudes across repetitions were obtained by taking mean of the average normalized waveform from all spikes in the transient window of each repetition. A scaling to range normalization method was performed as below.

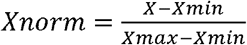

## Acknowledgements

The authors would like to acknowledge the members of the Pluta lab, Julia Veit, Kate Hong, Edward Bartlett and Daniel Butts for providing valuable feedback on the manuscript.

