## Supplementary Figure 1,2,3 & 4 for "Neural mechanisms for the localization of externally generated tactile motion"

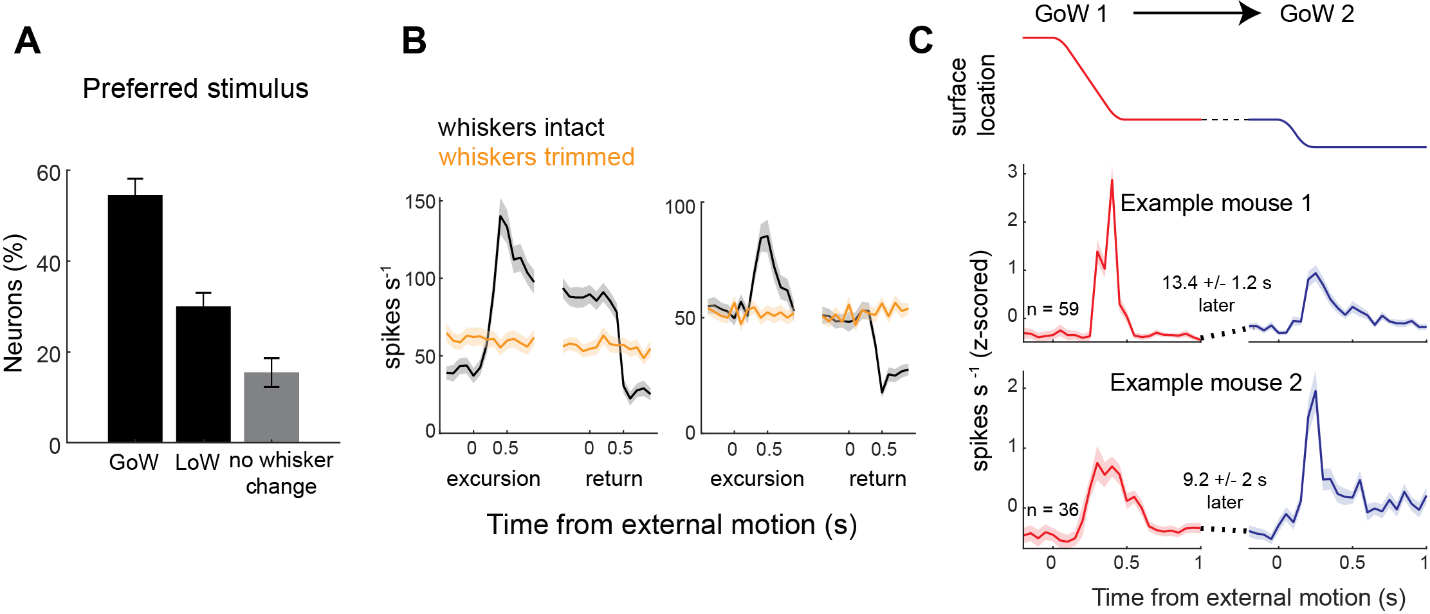

**Supplemental Figure 1. Neuronal firing rates are most responsive to gains in whisker contact**

**A)** Stimulus preference of neurons to gain of whisker, loss of whisker or no change in whiskers (385 neurons from 8 mice). **B)** The trial-averaged firing rate of two example neurons with intact whiskers and after all the whiskers were trimmed off. **C)** Population firing rates in 2 mice for trials where two GoWs occured successively (2 mice, 59 neurons and 36 neurons). GoW 1 and GoW 2 gained contact with different whiskers.

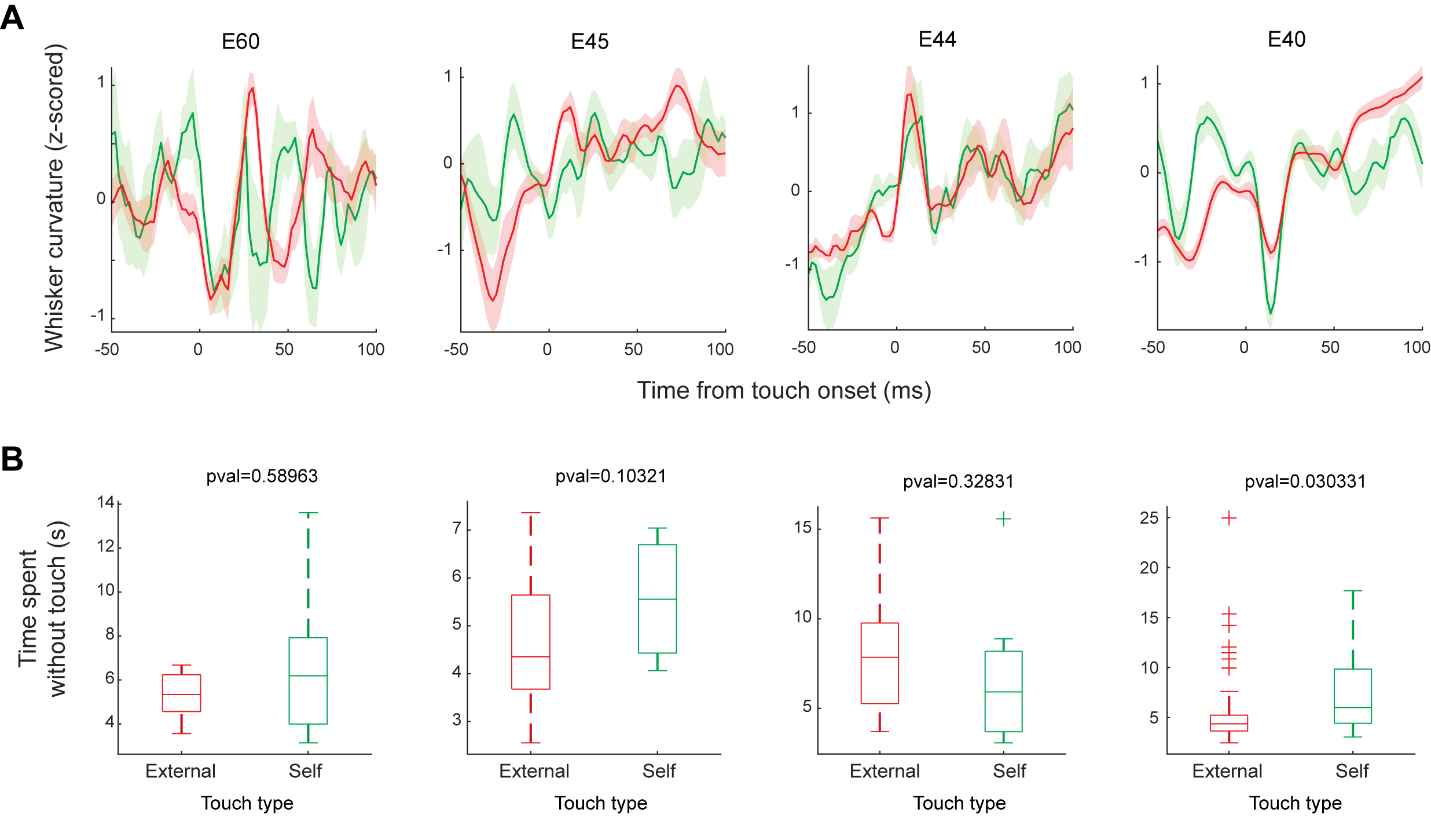

**Supplemental Figure 2. Whisker curvature and repetition rate were similar between external and self-generated touch events**

**A)** Trial-averaged whisker curvature aligned to the onset of touch during external and self-generated touch events. Each plot is from a separate animal (four total) **B)** The amount of time each animal spent without touch before experiencing an external or self-generated touch event. Each plot is from a separate animal (four total).

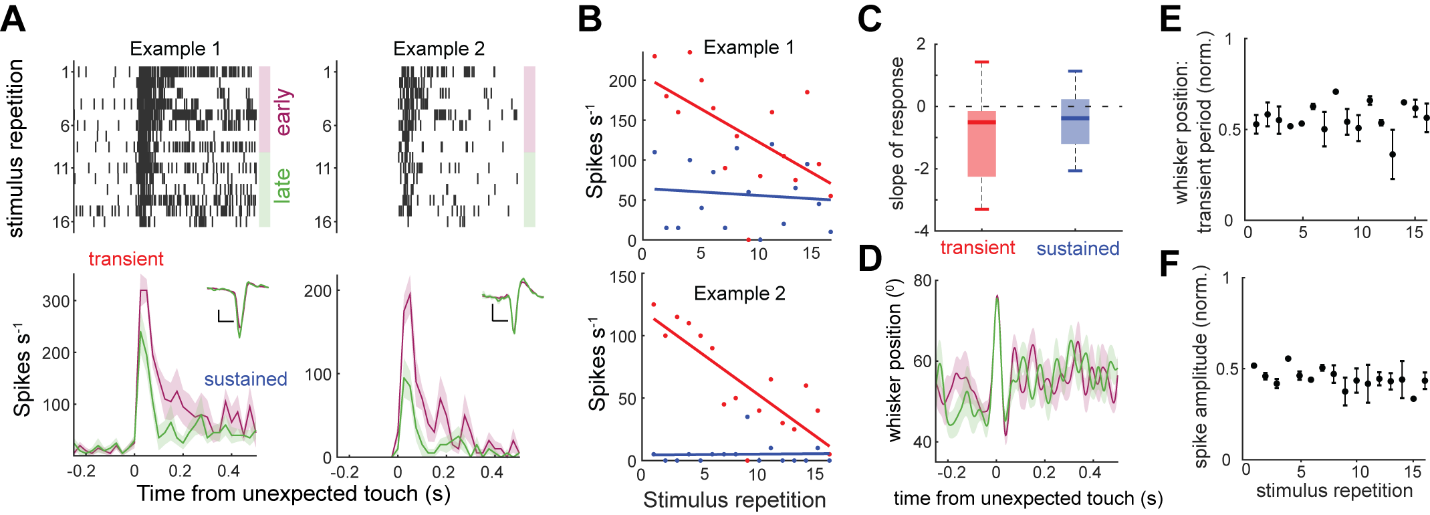

**Supplemental figure 3**. **Habituation of the transient response to unexpected touch**

**A)** Rasters and histograms of spike rate in two example SC neurons around the onset of unexpected touch. The histograms are divided by early (1^st^ half) and late (2^nd^ half) trials. **B)** Magnitude of the transient and sustained responses in these neurons as a function of stimulus repetition. Firing rate as a function of stimulus repetition was fit with a linear regression. **C)** Slope of the linear regression for each response type (40 transient responses, p < 0.001;14 sustained responses, p = 0.08; 2 mice). **D)** Comparison of whisker position for early (purple) and late (green) trials around the onset of unexpected touch. **E)** Normalized whisker position as a function of stimulus repetition. **F)** Normalized spike waveform amplitude as a function of stimulus repetition.

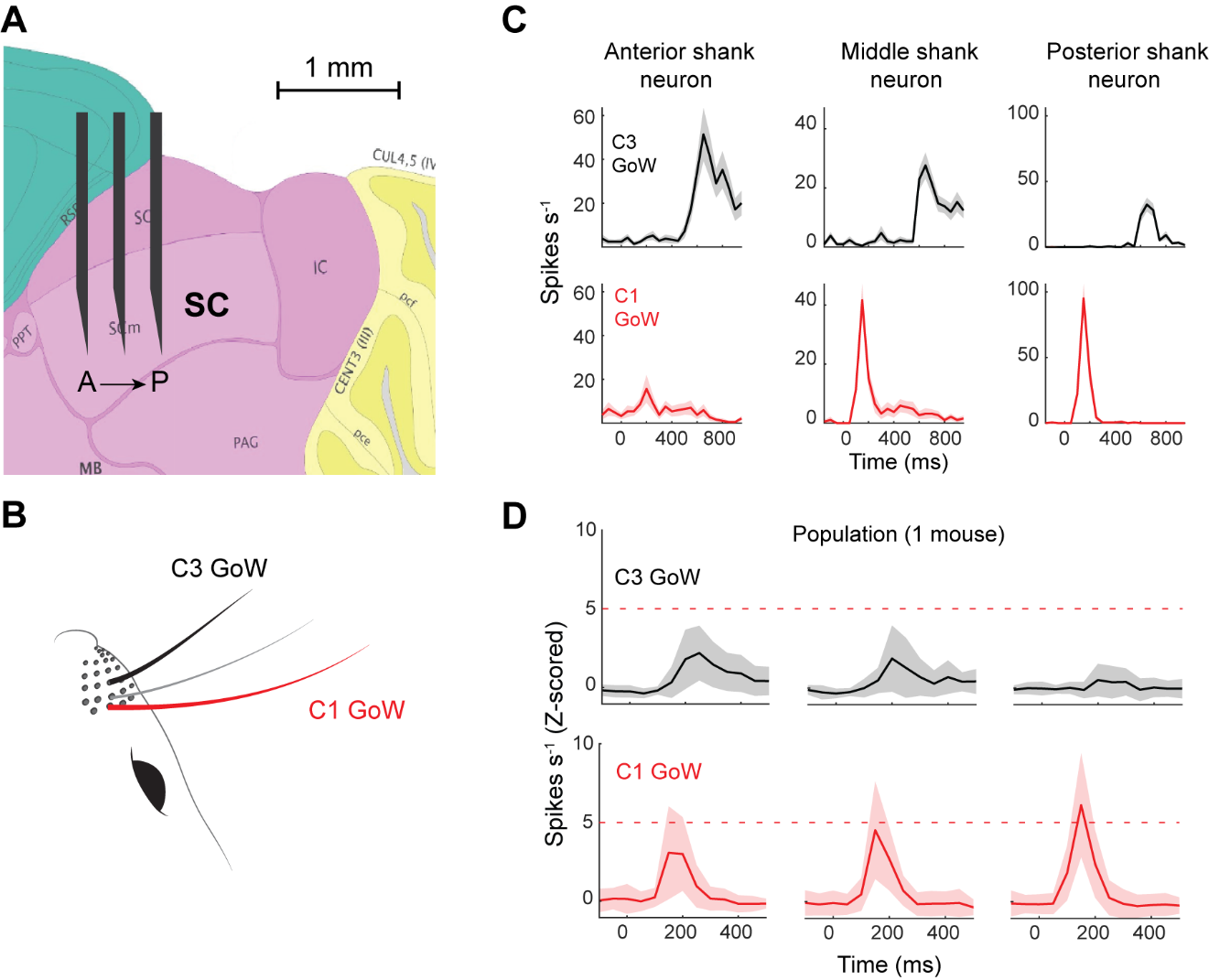
**Supplemental figure 4. The somatotopic organization of whisker space**

**A)** Orientation of electrode shanks in the SC. **B)** Illustration of whisker stimulation that generates neural responses in panel C and D. C1 GoW stimulation is represented in red, C3 GoW stimulation is in black. **C)** One example neuron from each shank arranged from anterior to posterior in the SC. **D)** Population averaged firing rates of neurons from 3 shanks (30, 23, 43 neurons per shank, from anterior to posterior, 1 mouse)
